# GAIA: an integrated metagenomics suite

**DOI:** 10.1101/804690

**Authors:** A. Paytuví, E. Battista, F. Scippacercola, R. Aiese Cigliano, W. Sanseverino

## Abstract

Identifying the biological diversity of a microbial population is of fundamental importance due to its implications in industrial processes, environmental studies and clinical applications. Today, there is still an outstanding need to develop new, easy-to-use bioinformatics tools to analyze both amplicon and shotgun metagenomics, including both prokaryotic and eukaryotic organisms, with the highest accuracy and the lowest running time. With the aim of overcoming this need, we introduce GAIA, an online software solution that has been designed to provide users with the maximum information whether it be 16S, 18S, ITS, or shotgun analysis. GAIA is able to obtain a comprehensive and detailed overview at any taxonomic level of microbiomes of different origins: human (e.g. stomach or skin), agricultural and environmental (e.g. land, water or organic waste). By using recently published benchmark datasets from shotgun and 16S experiments we compared GAIA against several available pipelines. Our results show that for shotgun metagenomics, GAIA obtained the highest F-measures at species level above all tested pipelines (CLARK, Kraken, LMAT, BlastMegan, DiamondMegan and NBC). For 16S metagenomics, GAIA also obtained excellent F-measures comparable to QIIME at family level. The overall objective of GAIA is to provide both the academic and industrial sectors with an integrated metagenomics suite that will allow to perform metagenomics data analysis easily, quickly and affordably with the highest accuracy.

## Introduction

The study of the different microbiomes present in either the environment or inside the human body is of fundamental importance as it is highly relevant for industrial processes as well as environmental and clinical applications. For instance, dysbiosis in distinct communities has been related to diseases or have been established as an early sign of environmental pollution [1-3].

Traditionally, studying bacterial communities required the isolation and culture of each individual microorganism, which is a significant limitation considering that less than 1% of the prokaryotes known are culturable [4]. Nowadays using high-throughput sequencing technologies is a standard for metagenomics analysis. These technologies are essential for the development of metagenomics, which is defined as the culture-independent genomic analysis of all the microorganisms in a particular environmental niche [5]. The data analysis can be either taxonomical, which aims to identify the origin of all the genomic material in the sample, or functional, where the goal is to identify the genes within a sample and their function in terms of Gene Ontology (GO) and metabolic pathways.

There are two main approaches in metagenomics: amplicon and shotgun analysis. Amplicon metagenomics is based on the PCR amplification of a genetic marker: the 16S rRNA in bacteria and archaea, ITS regions in fungi or the 18S rRNA in other eukaryotes. Even though it is a cheap and well-established technology, these markers lacks resolution as they cannot differentiate between closely related species [6]. Shotgun analysis is defined as the unrestricted sequencing of the genomes (Whole Genome Sequencing-WGS-metagenomics) or transcriptomes (metatranscriptomics) inside a sample. Shotgun analyses allow taxonomic identification down to strain level and also functional annotation by identifying genes present in the samples. In addition, the expression of these genes can be quantified by means of metatranscriptomics, thus allowing the identification of key roles of microorganisms in the samples. On the other hand, shotgun analyses are more computationally expensive then amplicon sequencing as they require higher sequencing coverage of the genomes in the sample. Additionally, the data analysis is more complex due to the large amount of data generated [7].

Dozens of pipelines have been developed so far for the analysis of metagenomics samples for taxonomic identification. Some of these pipelines have been compared in recent publications: Siegwald *et al*. (2017) [8] benchmarked commonly-used 16S pipelines, McIntyre *et al*. (2017) [9] benchmarked commonly-used shotgun metagenomics pipelines, and Brown, *et al*. (2017) [10] benchmarked pipelines that are able to process Nanopore data.

All things considered, there are still limitations to be overcome in the field of metagenomics: 1) there is still room for improvement in terms of accuracy, 2) some of the pipelines work on amplicon but not on shotgun analyses and vice versa, 3) lack of true user-friendly interfaces for the vast majority of the pipelines, 4) lack of pipelines that have eukaryotic databases, 5) high computational and data storage demand, especially in terms of shotgun metagenomics, and 6) high processing time. To overcome these limitations, we introduce GAIA: a new online metagenomics integrated suite which aims to become the reference method in metagenomics analysis for amplicon and WGS metagenomics as well as metatranscriptomics. With the aim of validating GAIA, we have gathered the results and datasets available from the different benchmarks, which include the following pipelines: BMP [11], mothur [12], QIIME [13], LMAT [14], BlastMegan [15], DiamondMegan [15], NBC [16], CLARK [17] and Kraken [18].

## Materials and methods

### Benchmark

#### Whole genome sequencing

The datasets (FASTA and FASTQ files) from Segata *et al*. (2013) and Ounit and Lonardi (2016) used in McIntyre, *et al*. (2017) for benchmarking were downloaded from http://ftp-private.ncbi.nlm.nih.gov/nist-immsa/IMMSA/. FASTA files were converted to FASTQ files using an in-house script and the quality assigned to the bases was 40 (Phred+33). The datasets were then mapped against the GAIA’s Prokaryotes database v1.0. Precision, recall and F-measure values at read-level for the benchmarked tools were extracted directly from the paper. Precision, recall and F-measures for GAIA were calculated as:

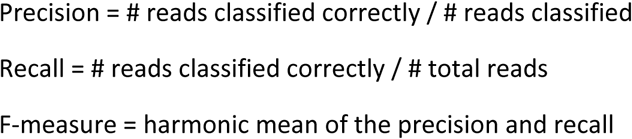

The datasets in FASTQ format from Brown, *et al*. (2017) were downloaded from the European Nucleotide Archive (ENA) as using the accessions PRJEB8672 and PRJEB8716. Accuracy values at read-level for the benchmarked tools were extracted directly from the paper. Accuracy for GAIA was calculated as:

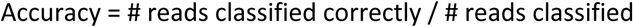

#### Amplicon sequencing

The datasets from Siegwald, *et al*. (2017) were downloaded from http://pegase-biosciences.com/metagenetics/. These datasets were then mapped against GAIA’s Amplicon NCBI 16s database v1.0. Precision, recall and F-measures at read-level for the benchmarked tools were extracted directly from the paper. Precision, recall and F-measure values for GAIA were calculated as:

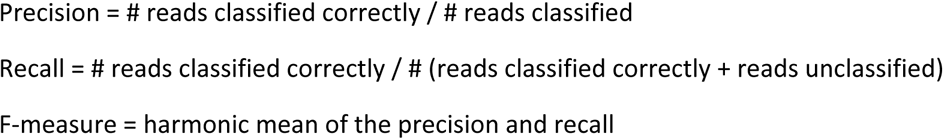

## Results

### Pipeline

GAIA consists of different Python, Java, Bash and R scripts that perform the following 5 different steps: i) the first step is the quality check and trimming, in that GAIA calls BBDuk [19] in order to remove both adapter sequences and bad quality portions from the reads for Illumina and Ion Torrent data and, for Oxford Nanopore data, it uses Porechop [20] for adapter removal; ii) BWA [21] is used to map the high quality reads from any platform (Illumina, Ion Torrent and Oxford Nanopore) against custom-made databases created from NCBI sequences [22]; iii) reads are classified into the most specific taxonomic level using an in-house Lowest Common Ancestor (LCA) algorithm; iv) minimum identity thresholds are applied to classify reads into strains, species, genus, family, order, class, phylum and domain levels; v) alpha and beta diversities are finally calculated using phyloseq [23]. Additionally, should any of the input datasets come from different conditions, GAIA includes an additional step to perform a differential abundance analysis using DESeq2 [24] (Figure 1).

**Figure 1.**
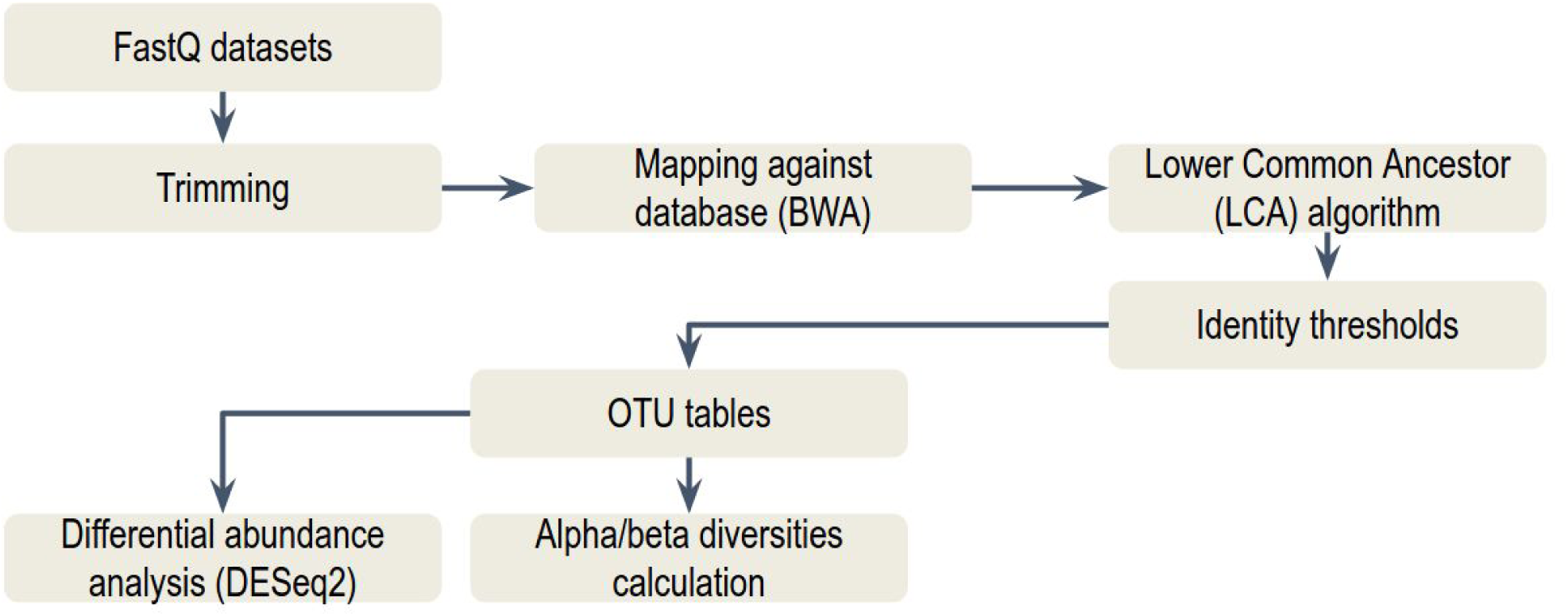
Overview of GAIA pipeline with the distinct steps it follows.

### Benchmark

In order to assess the performance of GAIA and to compare it with other metagenomics pipelines, we conducted a benchmark using both whole genome and amplicon sequencing data as described in the following paragraphs. For each dataset, GAIA’s precision, recall and F-score were calculated and compared with the others pipelines.

#### Whole genome sequencing

The datasets used from McIntyre, *et al*. (2017) were generated *in silico* and can be divided into two groups. The first group of datasets are created taking into account their complexity: high complexity datasets contain more species (100 different species) with more variable abundances than the low complexity datasets (25 different species). The second group of datasets were created including the species that are commonly found in mouth, city parks, gut or indoors. By comparing the results of GAIA with the expected ones, precision, recall and F-scores for each dataset were calculated (Supplementary Table 1). On average, GAIA obtained a precision of 0.982, a recall of 0.902 and the highest average F-score (0.94) at the species level, followed by CLARK-S and CLARK with F-scores of 0.936 and 0.921, respectively.

The Oxford Nanopore datasets used from Brown, *et al*. (2017) were real data generated from four separate cultures of *Escherichia coli, Pseudomonas fluorescens, Microcystis aeruginosa* and *Synechococcus elongatus*, and three mixed cultures of these four species. By comparing the results of GAIA with the expected ones, the accuracy for each dataset was calculated (Supplementary Table 2). On average, GAIA obtained the highest accuracy of 0.967, followed by Kraken with an accuracy of 0.946.

#### Amplicon sequencing

The datasets used from Siegwald, *et al*. (2017) were generated *in silico* with and without simulating sequencing errors on different rRNA subunits (V3, V4-V5) and can be divided into three groups:

- High Complexity (HC): all taxa equally distributed with no dominant organisms.
- Medium Complexity (MC): four dominant species of different genera accounting for 20% of reads and the remaining taxa are equally distributed.
- Low Complexity (LC): 1 dominant species accounting for 30% of reads and the remaining taxa are equally distributed.

At family level, using the Siegwald, *et al*. (2017) benchmark, GAIA obtained equal or slightly higher F-measures relative to QIIME: GAIA (0.957), QIIME UCLUST with SILVA database (0.956), QIIME SortMeRna SUMACLUST with Greengenes database (0.955) (Supplementary Table 3). At genus level, GAIA (0.83) showed the third best F-measure after CLARK (0.878) and Kraken (0.859), which was followed by QIIME UCLUST with SILVA database (0.776) and QIIME SortMeRna SUMACLUST with Greengenes database (0.665) (Supplementary Table 4).

### Online platform

In order to provide a comfortable user experience, the GAIA pipeline was integrated into an online software solution, which delivers the software in a way that can be accessed from any device with an Internet connection and a web browser without any bioinformatics skills required. The analysis is performed interactively online and it includes dynamic charts and tables using Google Charts and DataTables (JavaScript-based) (Figure 2).

**Figure 2.**
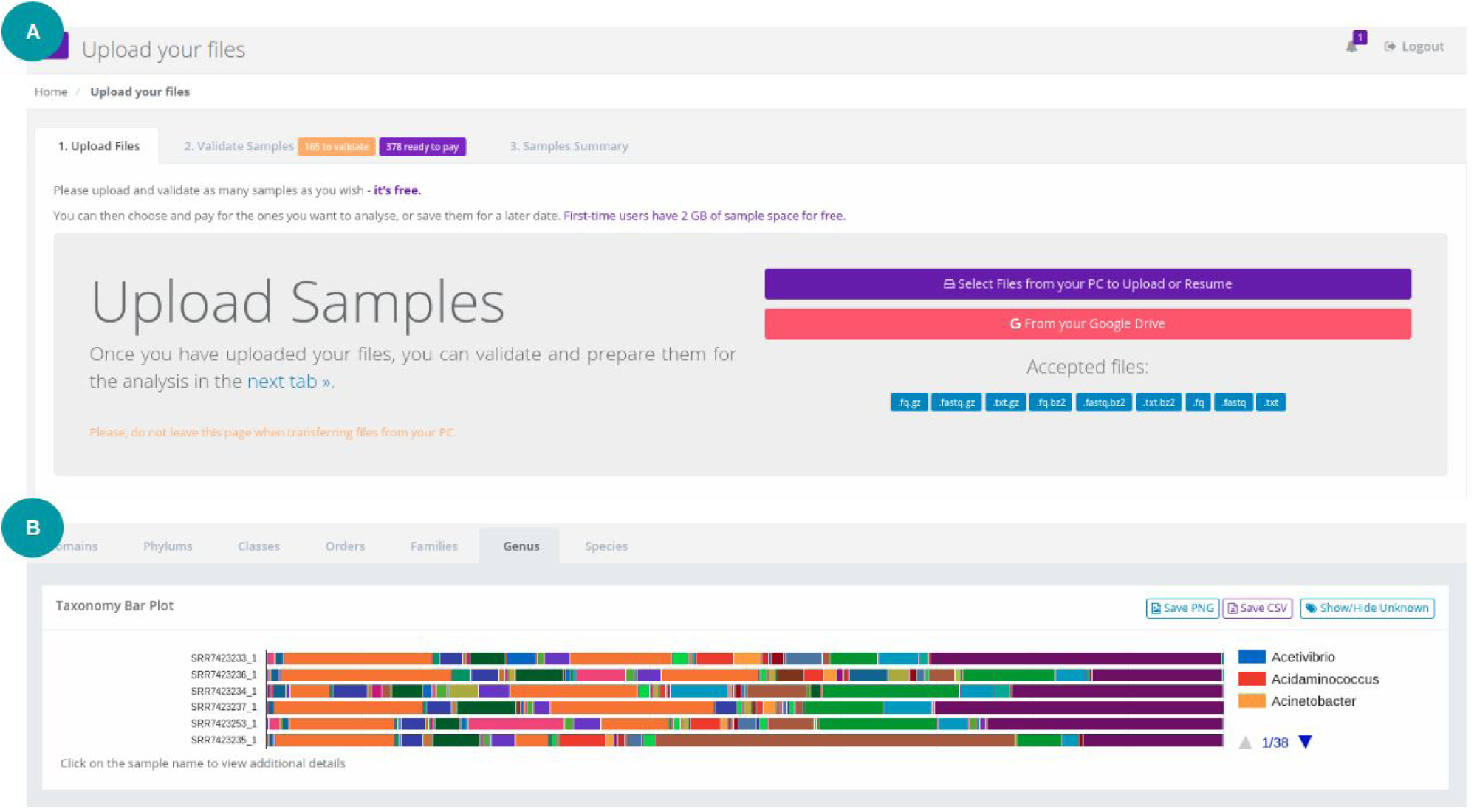
Screenshot of the upload page (A), in which the user uploads sequencing data, and screenshot of the taxonomy barplot at genus level once the analysis has been completed (B).

## Conclusions

We propose GAIA as a new software able to obtain a comprehensive and detailed overview at any taxonomic level (including strains) of microbiomes of different origins such as human (e.g. stomach or skin), agricultural and environmental (e.g. land, water or organic waste) in an accurate and easy way. The presented high benchmark scores validate the algorithm. In fact, on average GAIA obtained the highest scores at species level for WGS metagenomics and it also obtained excellent scores for amplicon sequencing. Indeed, GAIA was the most performing software at genus level and within the top-three most performing softwares at family level for 16S rRNA data. In addition, at both family and genus level for amplicon sequencing, GAIA obtained higher F-scores than QIIME, the most cited software for this kind of analysis. As metagenomics is shifting towards shotgun analyses which are able to sequence any organisms within a sample, GAIA’s database also includes eukaryotes to perform the so-called true metagenomics: a complete view in terms of existing life within the samples. GAIA has been created so the user can spend more time interacting with their results and less time setting up the analysis. The overall objective of GAIA is to provide academia and industries with an integrated metagenomics suite that will allow to perform metagenomics data analysis easily and quickly. GAIA is available at http://gaia.sequentiabiotech.com.

## Acknowledgements

We thank Anna Delgado Tejedor for her contribution in evaluating GAIA during her internship at Sequentia Biotech.

## References

[1] Lynch, S. V., & Pedersen, O. (2016). The Human Intestinal Microbiome in Health and Disease. The New England Journal of Medicine, 375(24), 2369–2379. https://doi.org/10.1056/NEJMra1600266=

[2] McLellan, S. L., Fisher, J. C., & Newton, R. J. (2015). The microbiome of urban waters. International Microbiology, 18(3), 141–149. https://doi.org/10.2436/20.1501.01.244=

[3] Rajagopala, S. V., Vashee, S., Oldfield, L. M., Suzuki, Y., Venter, J. C., Telenti, A., & Nelson, K. E. (2017). The human microbiome and cancer. Cancer Prevention Research, 10(4), 226–234. https://doi.org/10.1158/1940-6207.CAPR-16-0249=

[4] Schloss, P. D., & Handelsman, J. (2005). Metagenomics for studying unculturable microorganisms: cutting the Gordian knot. Genome Biology, 6(8), 229. https://doi.org/10.1186/gb-2005-6-8-229=

[5] Goodwin, S., McPherson, J. D., & McCombie, W. R. (2016). Coming of age: ten years of next-generation sequencing technologies. Nature Reviews Genetics, 17(6), 333–351. https://doi.org/10.1038/nrg.2016.49=

[6] Perrine, H., Lagier, J.-C., Colson, P., Bittar, F., & Raoult, D. (2017). Repertoire of human gut microbes. Microbial Pathogenesis, 106, 103–112. https://doi.org/10.1016/J.MICPATH.2016.06.020=

[7] Ranjan, R., Rani, A., Metwally, A., McGee, H. S., & Perkins, D. L. (2016). Analysis of the microbiome: Advantages of whole genome shotgun versus 16S amplicon sequencing. Biochemical and Biophysical Research Communications (Vol. 469). Elsevier Ltd. https://doi.org/10.1016/j.bbrc.2015.12.083=

[8] Siegwald, L., Touzet, H., Lemoine, Y., Hot, D., Audebert, C., & Caboche, S. (2017). Assessment of Common and Emerging Bioinformatics Pipelines for Targeted Metagenomics. PloS One, 12(1), e0169563. https://doi.org/10.1371/journal.pone.0169563=

[9] McIntyre, A. B. R., Ounit, R., Afshinnekoo, E., Prill, R. J., Hénaff, E., Alexander, N., Mason, C. E. (2017). Comprehensive benchmarking and ensemble approaches for metagenomic classifiers. Genome Biology, 18(1), 182. https://doi.org/10.1186/s13059-017-1299-7=

[10] Brown, B. L., Watson, M., Minot, S. S., Rivera, M. C., & Franklin, R. B. (2017). MinION™ nanopore sequencing of environmental metagenomes: a synthetic approach. GigaScience 6(3), 1–10. http://doi.org/10.1093/gigascience/gix007=

[11] Pylro VS, Roesch LFW, Morais DK, Clark IM, Hirsch PR, Tótola MR. Data Analysis for 16S Microbial Profiling from Different Benchtop Sequencing Platforms. J Microbiol Methods 2014;107:30–7. pmid:25193439

[12] Schloss, P. D., Westcott, S. L., Ryabin, T., Hall, J. R., Hartmann, M., Hollister, E. B., et al. (2009). Introducing mothur: open-source, platform-independent, community-supported software for describing and comparing microbial communities. Applied and Environmental Microbiology 75, 7537–7541. http://doi.org/10.1128/AEM.01541-09=

[13] Caporaso, J. G., Kuczynski, J., Stombaugh, J., Bittinger, K., Bushman, F. D., Costello, E. K., et al. (2010). QIIME allows analysis of high-throughput community sequencing data. Nature Methods 7, 335–336. https://doi.org/10.1038/nmeth.f.303=

[14] Ames, S. K., Hysom, D. A., Gardner, S. N., Lloyd, G. S., Gokhale, M. B., Allen, J. E. (2013). Scalable metagenomic taxonomy classification using a reference genome database. Bioinformatics (Oxford, England) 29(18), 2253–2260. http://doi.org/10.1093/bioinformatics/btt389=

[15] Huson, D. H., Auch, A. F., Qi, J., Schuster, S. C. (2007). MEGAN analysis of metagenomic data. Genome research 17(3), 377–386. http://doi.org/10.1101/gr.5969107=

[16] Rosen, G. L., Reichenberger, E. R., Rosenfeld, A. M. (2011). NBC: the Naive Bayes Classification tool webserver for taxonomic classification of metagenomic reads. Bioinformatics (Oxford, England) 27(1), 127–129. http://doi.org/10.1093/bioinformatics/btq619=

[17] Ounit, R., Wanamaker, S., Close, T. J., Lonardi, S. (2015). CLARK: fast and accurate classification of metagenomic and genomic sequences using discriminative k-mers. BMC Genomics 16, 236. https://doi.org/10.1186/s12864-015-1419-2=

[18] Wood, D. E., Salzberg, S. L. (2014). Kraken: ultrafast metagenomic sequence classification using exact alignments. Genome Biology 15, R46. https://doi.org/10.1186/gb-2014-15-3-r46=

[19] BBDuk Guide - DOE Joint Genome Institute. (n.d.). Retrieved February 18, 2018, from https://jgi.doe.gov/data-and-tools/bbtools/bb-tools-user-guide/bbduk-guide/=

[20] Wick, R. (2017). Adapter trimmer for Oxford Nanopore reads. Available online at: https://github.com/rrwick/Porechop=

[21] Li, H., & Durbin, R. (2010). Fast and accurate long-read alignment with Burrows–Wheeler transform. BIOINFORMATICS ORIGINAL PAPER, 26(5), 589–59510. https://doi.org/10.1093/bioinformatics/btp698=

[22] Database resources of the National Center for Biotechnology Information NCBI Resource Coordinators. (2016). Nucleic Acids Research, https://doi.org/10.1093/nar/gkv1290=

[23] Mcmurdie, P. J., Holmes, S., & Watson, M. (2013). phyloseq: An R Package for Reproducible Interactive Analysis and Graphics of Microbiome Census Data. PLoS ONE, 8(4). https://doi.org/10.1371/journal.pone.0061217=

[24] Love, M. I., Huber, W., & Anders, S. (2014). Moderated estimation of fold change and dispersion for RNA-seq data with DESeq2. Genome Biology, 15. https://doi.org/10.1186/s13059-014-0550-8=

